# Graph neural network based heterogeneous propagation scheme for classifying alzheimer’s disease

**DOI:** 10.1101/2021.01.21.427712

**Authors:** Jiyoung Byun, Yong Jeong

**Affiliations:** Department of Bio and Brain Engineering Korea Advanced Institute of Science and Technology, Daejeon 34141, Republic of Korea

**Keywords:** Alzheimer’s disease, Classification, Resting state functional connectivity, Deep learning, Graph neural network

## Abstract

Deep learning frameworks for disease classification using neuroimaging and non-imaging information require the capability of capturing individual features as well as associative information among subjects. Graphs represent the interactions among nodes, which contain the individual features, through the edges in order to incorporate the inter-relatedness among heterogeneous data. Previous graph-based approaches for disease classification have focused on the similarities among subjects by establishing customized functions or solely based on imaging features. The purpose of this paper is to propose a novel graph-based deep learning architecture for classifying Alzheimer’s disease (AD) by combining the resting-state functional magnetic resonance imaging and demographic measures without defining any study-specific function. We used the neuroimaging data from the ADNI and OASIS databases to test the robustness of our proposed model. We combined imaging-based and non-imaging information of individuals by categorizing them into distinctive nodes to construct a *subject–demographic bipartite graph*. The approximate personalized propagation of neural predictions, a recently developed graph neural network model, was used to classify the AD continuum from cognitively unimpaired individuals. The results showed that our model successfully captures the heterogeneous relations among subjects and improves the quality of classification when compared with other classical and deep learning models, thus outperforming the other models.

## 1 Introduction

Alzheimer’s disease (AD) is the most common type of dementia worldwide, and is characterized by initial memory impairment and additional cognitive dysfunction as the disease progresses [1]. In the past, the diagnosis of AD was based on eliminating the possibility of other potential diseases rather than on pathological proof. Moreover, AD is usually identified with other types of neurodegenerative diseases; thus, the definitive confirmation of clinical AD was not available unless neuropathology was evaluated post-mortem. Clinical AD diagnosis is even harder during a long period of preclinical asymptomatic phase because cognitively unimpaired individuals can also have the disease [2].

Although AD is becoming more common in an aging society, developing treatments for it is challenging. The rate of failure in drug development is high and, therefore, no new drugs have been approved since 2003 and no disease-modifying treatments have been approved [3]. Therefore, early risk-reducing interventions based on biomarkers to delay the progression or to prevent the onset of AD are regarded as important issues [4]. Neuroimaging biomarkers, such as structural magnetic resonance imaging (sMRI), positron emission tomography (PET), and functional MRI (fMRI) have been developed and used clinically to identify AD. These approaches have advantages over fluid-based approaches because they are non-invasive and can be used to distinguish the different phases of AD [5].

fMRI is associated with neuronal activity and detects the blood-oxygenation-level-dependent (BOLD) signal. Although it can be examined with or without the application of a task, the latter strategy, which is also called resting-state fMRI (rs-fMRI), has advantages in clinical conditions [6, 7, 8, 9]. Task-fMRI is difficult to apply to cognitively impaired individuals because it often requires a longer time than rs-fMRI for data acquisition, and the result highly depends on task performance. On the other hand, rs-fMRI offers a better signal-to-noise ratio while efficiently representing brain metabolism with a relatively simple data acquisition process. It has also been shown that abnormal functional connectivity assessed using rs-fMRI is related to regions of early brain atrophy and hypometabolism in AD [10, 11, 12, 13, 14].

Progress in neuroimaging technologies has accelerated the development of computer-aided approaches to produce reliable diagnosis from large-scale and high-dimensional brain imaging data. Techniques such as support vector machines, linear discriminant analysis, and linear regression have been used for AD classification [15]. The deep learning approach, which automatically learns discriminative features through a hierarchical learning process, requires little or no pre-processing steps and can select features in an end-to-end manner without human intervention [16]. All steps are simultaneously optimized, which, in turn, leads to enhanced performance by detecting unbiased features from complex data. Therefore, it has been successfully applied to AD classification using several different imaging modalities, such as sMRI [17], PET [18, 19], and rs-fMRI [20].

Graph neural network (GNN) was introduced to efficiently process data that are presented in a graphical format [21, 22]. Graphs are composed of nodes and edges, which characterize objects and their relationships, respectively. In other words, assigning edges can represent the associations or similarities between nodes. It has been applied in various areas such as natural science (physics, chemistry, biology), social science, and finance. GNN was also applied in neuroimaging studies to predict autism spectrum disorder, brain tumor, and bipolar disorder. Ktena et al. (2018) classified the autism spectrum disorder and sex by granting structural and functional information to the edges in a graph [23]. Similarity between the nodes in the constructed graph was learned through the spectral graph convolutional network (GCN). Moreover, Parisot et al. (2018) tested the influence of the graph structure on spectral GCN when classifying autism spectrum disorder and Alzheimer’s disease [24].

Because GCN averages the information from neighboring nodes during the learning processes of similar properties among adjacent nodes, the ‘over-smoothing’ issue occurs during the propagation of multiple layers [25, 26]. Loss of useful information will prevent the model from considering higher-order information and eventually limits the performance. Moreover, previous studies applying graph-based classification approaches [27, 24] and *subject–subject graph* based on handcrafted patient similarity functions have been widely reported. Nevertheless, such graph construction methods have difficulties in expanding the number and types of subject features; it is not only difficult to manually determine the precise importance weight of each feature, but the scalar-valued similarity (i.e., the weight of an edge) restricts the representation power of the model. Furthermore, similarity functions cannot compute the weight if the dataset contains missing features, which commonly occurs in real-world applications.

To resolve the aforementioned limitations, we developed a graph-based deep learning framework for classifying the AD continuum. First, we constructed a *subject–demographics graph* to incorporate all the information. Because it is not necessary to use study-specific functions to incorporate distinct information, potential bias will be relieved. In addition, this approach can readily incorporate heterogeneous data, which increases its generalizability. Second, the implementation of approximate personalized propagation of neural predictions (APPNP) ameliorates the over-smoothing problem by introducing a propagation scheme of personalized PageRank [28]. Moreover, APPNP does not contain learnable parameters in the propagation stages, which in turn helps the model to avoid overfitting.

To evaluate the performance of the proposed method, it was implemented on the Alzheimer’s Disease Neuroimaging Initiative (ADNI) and Open Access Series of Imaging Studies (OASIS) datasets. We used functional connectivity from rs-fMRI and additional relevant information (e.g., age, sex, and APOE genotype) as the inputs in this work. Features from imaging and non-imaging data were automatically estimated using multilayer perceptron (MLP). This information was then propagated along the edges in the *subject–demographics graph*. The overall results show that our proposed formulation for integrating the features to construct a graph and learning through APPNP improves the classification performance.

## 2 Methods

### 2.1 Neuroimaging data

The first dataset was obtained from phase 2 of the ADNI dataset (http://adni.loni.usc.edu/). The aim of ADNI is to detect and track AD progression at the earlier stages with identified biomarkers based on MRI, PET, and clinical and neuropsychological assessment. In this study, we used sMRI (T1-weighted MPRAGE sequence) for preprocessing and rs-fMRI scans from the ADNI2 dataset for classification. Among subjects who were initially diagnosed as cognitively unimpaired (CU) or AD, we excluded those who did not undergo rs-fMRI scanning or the scan was of problematic quality. Thus, a total of 62 subjects were included (CU, n = 31; AD, n = 31).

To assess the applicability of our proposed model to different datasets, we also used the OASIS phase 3 dataset (https://www.oasis-brains.org) [29], which is a longitudinal clinical neuroimaging dataset for normal aging and AD. A total of randomly selected 240 subjects who were initially diagnosed as cognitively unimpaired (CU, n = 120) or AD (n = 120) were included. A detailed protocol for data acquisition can be found in the official webpage of each dataset.

### 2.2 Image preprocessing

Preprocessing of rs-fMRI was conducted using Statistical Parametric Mapping (SPM12; Wellcome Trust Centre for Neuroimaging, London, UK, http://www.fil.ion.ucl.ac.uk/spm/software/spm12) [30] in MATLAB 2019a.

The first 5 echo planar imaging volumes were discarded for signal stabilization. After obtaining the images of each participant, different signal acquisition times were corrected for the rest by shifting the signals of each slice to the midpoint of TR. The sMRI of each participant was co-registered to the mean fMRI to segment it into gray matter, white matter, and CSF. Nuisance covariate regression for 6 parameters was conducted to eliminate head motion effects, which are known to cause interference when measuring functional connectivity. The detrended time series was calculated by subtracting the linear trend. Spatially normalized images to the MNI template were smoothed at 6 mm FWHM. Lastly, all preprocessed rs-fMRI were band-pass filtered (0.01 -– 0.1 Hz.)

### 2.3 Brain functional connectivity construction

For constructing the brain functional connectivity, we used R = 300 network liberal mask as the region of interest (ROI). It is a widely used parcellation of the cerebral cortex based on the resting-state functional connectivity [31]. The preprocessed rs-fMRI time series was extracted and averaged in each ROI. We utilized the available function (i.e., correlation) in Nilearn [32], which is a Python module for efficient statistical learning of neuroimaging data. The raw values of the calculated resting state functional connectivity (RSFC) were provided as inputs to our proposed framework. Further information regarding each measure is available at the Nilearn official website (https://nilearn.github.io/#).

### 2.4 Graph neural network-based classification

This paper proposes a deep learning-based classification approach, which is composed of two stages: (1) feature extraction using MLP and (2) graph propagation using a GNN. In short, stage (1) takes the input of the RSFC matrix calculated from rs-fMRI and non-imaging demographic measures such as the APOE genotype, age, and sex. Based on the given raw features, the model extracts informative feature vectors, which are used for the initial node representation. Stage (2) integrates the extracted features and propagates the information along the edges in the *subject–demographics graph*. The obtained graph representation is utilized for disease classification via a fully connected sigmoid layer. The proposed architecture is fully differentiable, and it is trained in an end-to-end manner. Fig. 3 is the graphical illustration of our model.

**Figure 1:**
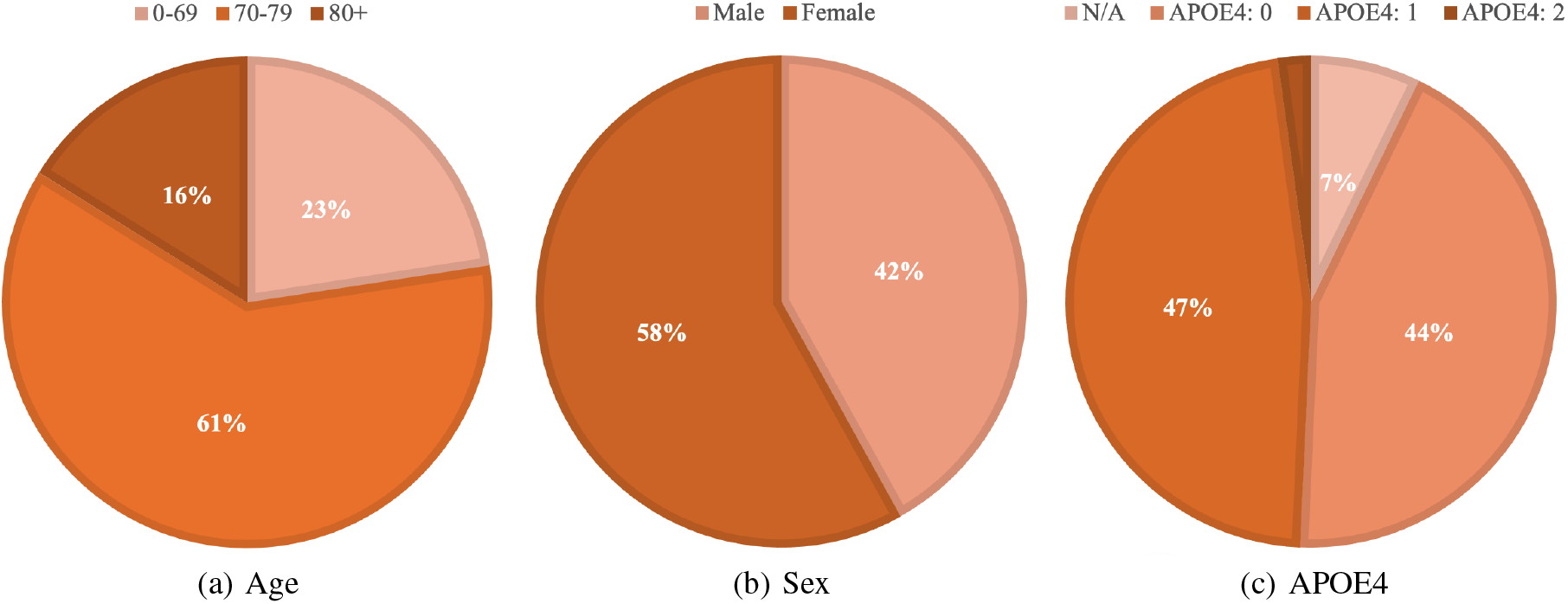
Ratio of each demographic measure in ADNI dataset.

**Figure 2:**
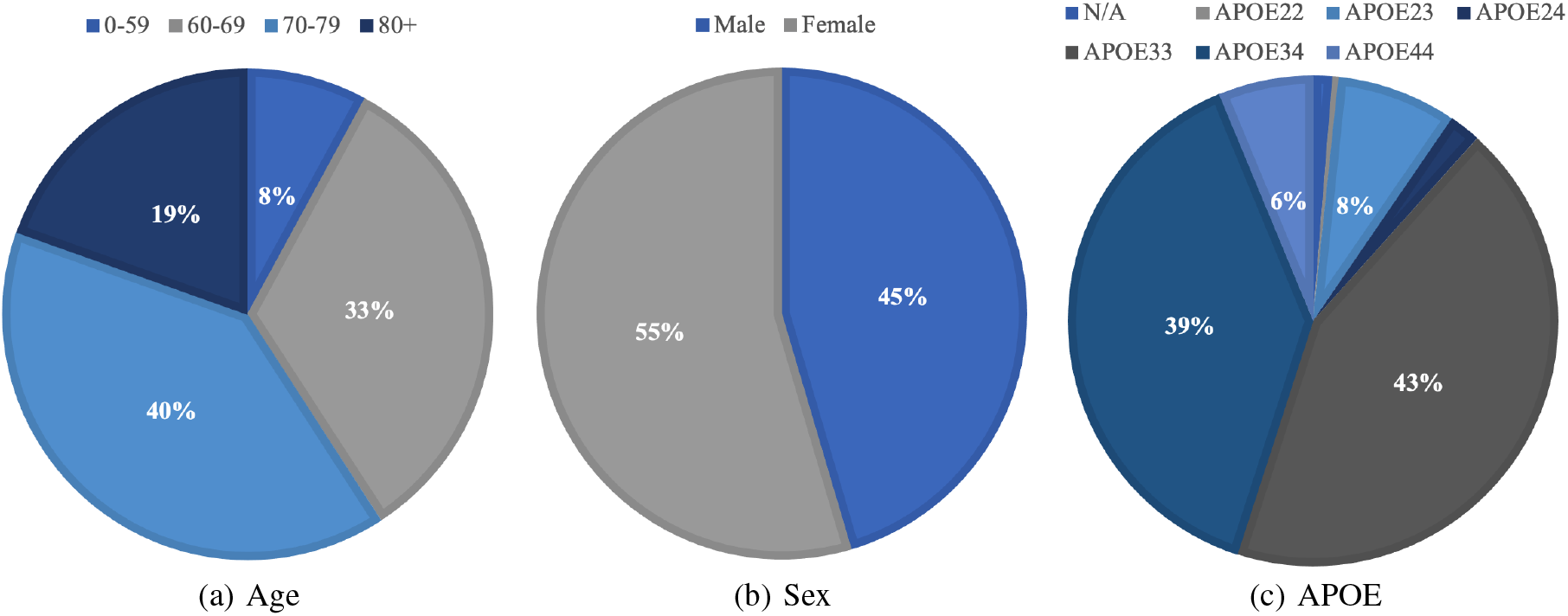
Ratio of each demographic measure in OASIS dataset.

**Figure 3:**
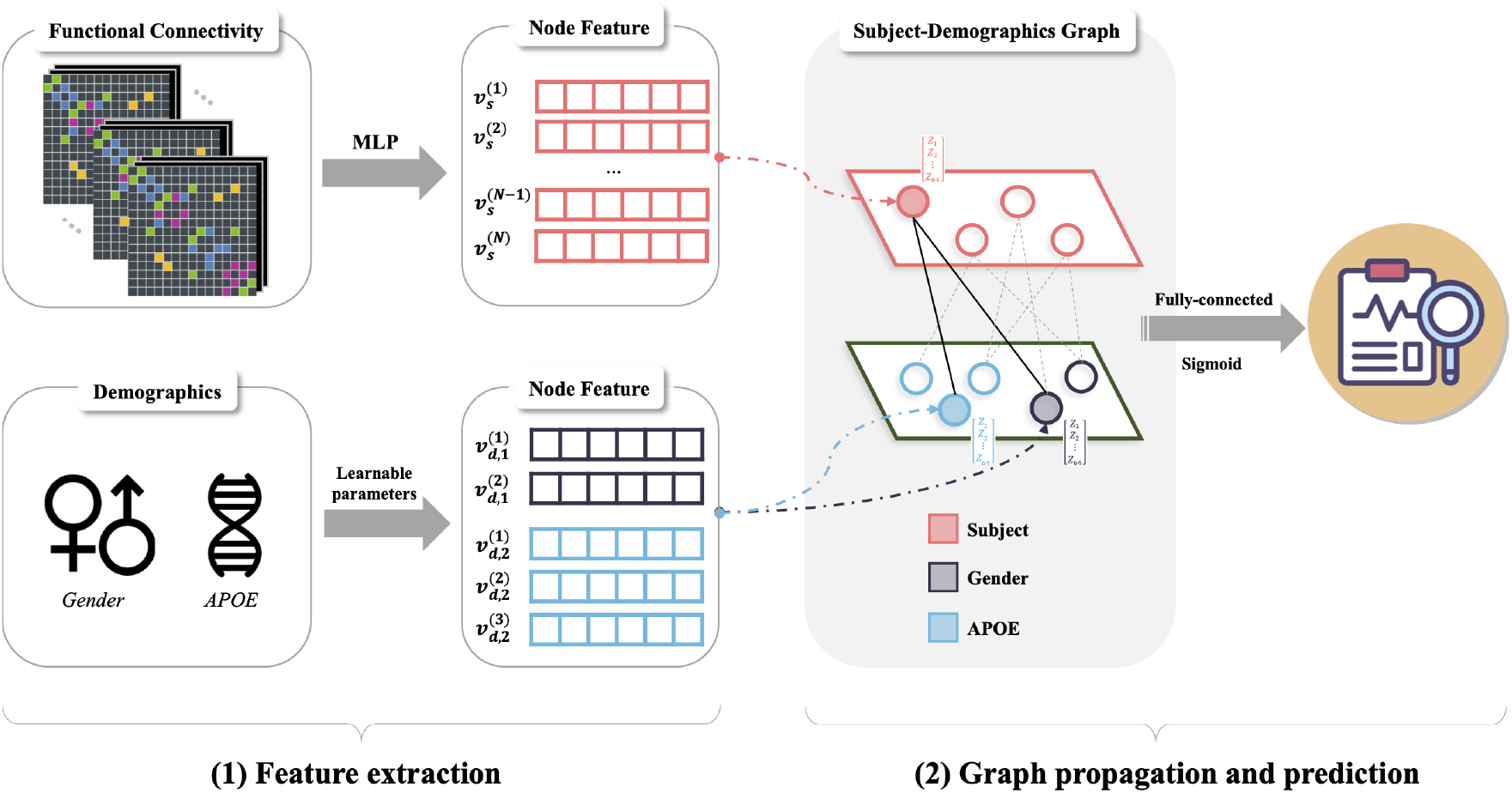
APPNP architecture of the proposed deep-learning based classification framework.

**Algorithm 1.**
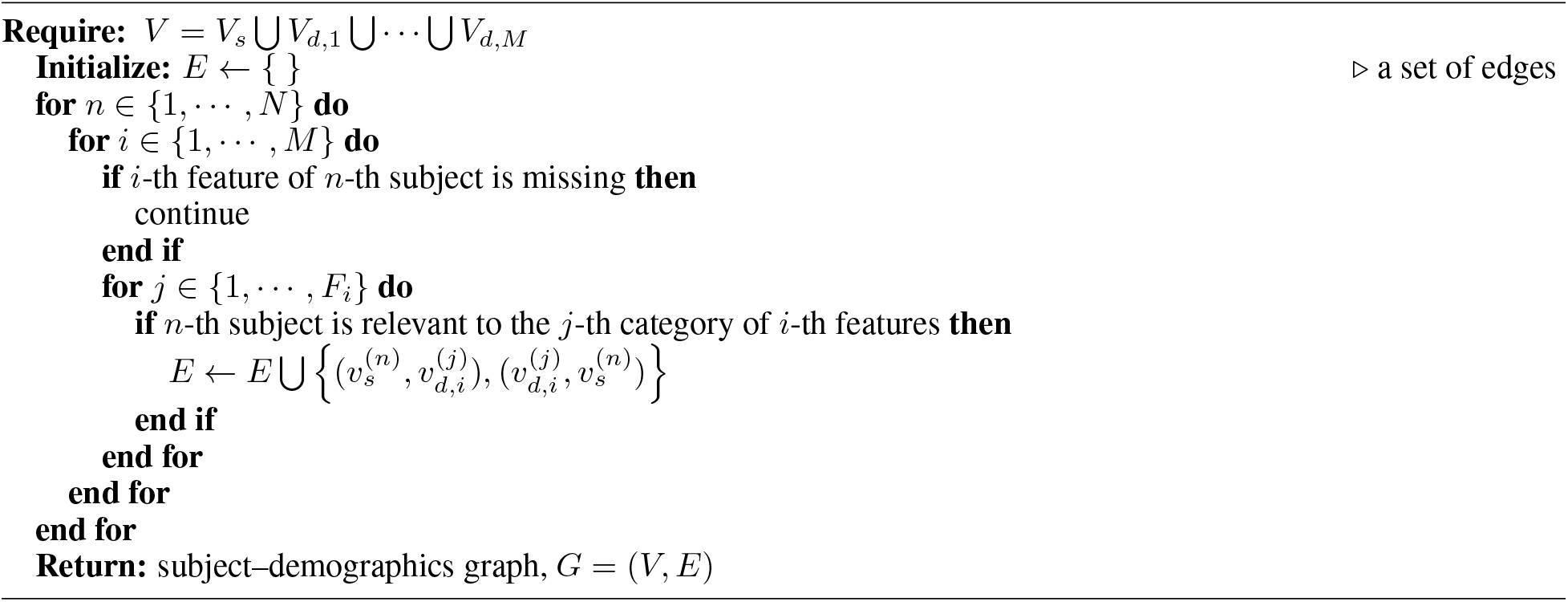
Subject-Demographics Graph Construction

#### 2.4.1 Subject-demographics graph construction

Considering a dataset with *N* subjects and *M* different categorical demographic features, we first define the graph *G*, which consists of a set of *N* subject nodes *V_s_* and *M* sets of demographic measure nodes *V_d_*:

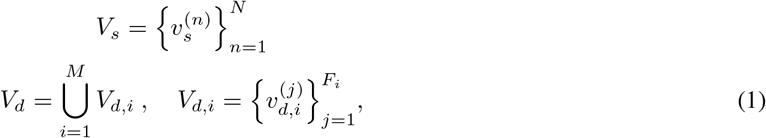

where 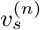 is a node of the *n*-th subject, 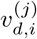 is a node of the *j*-th category of the *i*-th feature, and *F*_*i*_ is the cardinality of the *i*-th feature. For example, in the case of the gender feature (e.g., let *i*=1), the first and second gender nodes 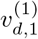 and 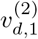 represent the male and female nodes, respectively, and the cardinality of gender is two (*F*_1_=2). The total number of nodes in the graph is 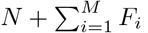. For each subject and demographic feature, we connected the corresponding subject node to the relevant feature node. We do not insert an edge for a missing demographic feature. The detailed graph construction process is provided in Algorithm 1. Because the components of the demographic features in the subject–demographics graph would influence the performance of the proposed model, we evaluated the classification accuracy depending on the demographic measures in section 3.3.

#### 2.4.2 Feature extraction

Following the idea of *representation learning*, which has achieved remarkable success in various artificial intelligence research areas [33, 34, 35, 36, 22], this study adopted a shallow MLP for extracting the latent representation of each patient node from the raw data (i.e., entries of the RSFC).

Primarily, we vectorized all entries of the RSFC for each subject and generated a matrix 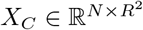. To complement the feature information, one-hot encoded categorical demographic features, 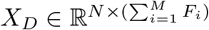, were also evaluated. The input matrix *X* was defined by concatenating two matrices horizontally: 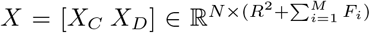

This network can be represented as

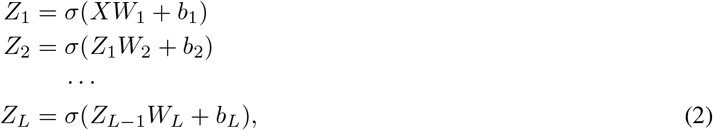

where *W_l_* is the weight matrix, *b_l_* is the bias vector for the *l*-th layer, and *σ*(·) is the activation function, which is ReLU in our model. The output of the last layer, *Z_L_*, denotes the *H*-dimensional feature vectors that function as input vectors for the graph neural network.

#### 2.4.3 Graph propagation

As demonstrated in section 2.4.1, our proposed GNN-based classification model is composed of two types of nodes: subject-specific and demographics nodes. First, the outputs of the MLP, *Z_L_*, are taken as input for the subject-specific nodes. Then, we allocate the vectorized values with the same dimension as that of the subject-specific nodes to the demographic nodes. These nodes are filled with learnable parameters updated in an end-to-end manner. To elaborate this, we denote *D* as the initial feature matrix for the demographic nodes. 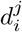, which represents the *j*-th category of the vectorized value of the *i*-th demographic feature node, is assigned to the 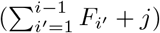-th row of *D*.

The majority of conventional graph neural network-based disease classification models have used the GCN. Despite some successes in applying this method, it has limitations related to the shallow learning mechanisms [33, 37]. This might result in the failure to discover complex patterns beyond the graphs, which affects the performance of the model. Therefore, we utilized the APPNP scheme for the learning process. This approach not only integrates the information from the neighboring nodes, it also maintains the locality by adjusting the teleport probability *α*. Consequently, this algorithm prevents the over-smoothing issue and allows deeper GNN models [38]. Moreover, APPNP does not have learnable parameters during the propagation steps; it also alleviates the overfitting issue. In section 3.2, we compared the accuracy of GCN and APPNP depending on the number of propagations (hops) on both the ADNI and OASIS datasets. The propagation steps in the APPNP scheme are summarized as follows:

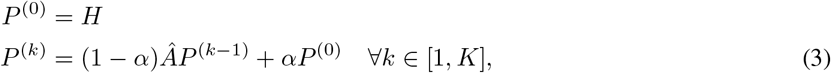

where *H* = [*Z_L_*; *D*] is a vertically concatenated matrix and *K* is defined as the number of propagation steps. 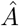 is a symmetrically normalized adjacency matrix:

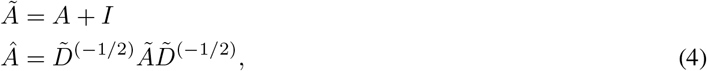

where *A* is an adjacency matrix of the graph *G*, as described in section 2.4.1 and Algorithm 1, and 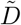 is a diagonal degree matrix of 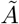.

#### 2.4.4 AD classification

Using the graph representation of the evaluated subject nodes in Eq. 3, we classified the *i*-th subject, 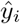, as CU or AD using a fully connected sigmoid layer as follows:

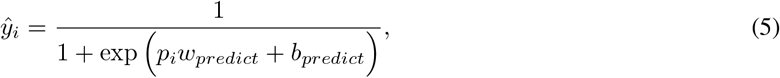

where *p_i_* ∈ ℝ^1*×H*^ is the *i*-th row of *P*^(*K*)^. *w_predict_* ∈ ℝ^*H×*1^ and *b* ∈ ℝ^1×1^ are learnable parameters in the fully connected layer. Finally, all learnable parameters in the proposed model were optimized to minimize the loss 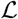 using the adaptive moment estimation (Adam) algorithm [39].

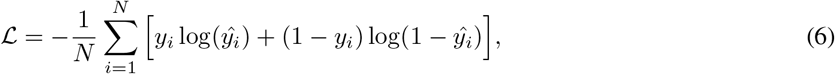

where *y_i_* = 1 when the *i*-th subject is an AD patient and *y_i_* = 0 otherwise. The deep-learning-based framework was implemented in Python using PyTorch [40], a library for machine learning and deep learning, on a single NVIDIA GPU.

## 3 Results

### 3.1 Experimental setup

To test the generalizability of the proposed model, conventional model assessment methods including *k*-fold cross validation (CV) were performed. Leave-one-out cross-validation (LOOCV) was less biased than *k*-fold CV. This finding can be attributed to the tendency of negligible overestimation of an error and difficulties in leaving out a substantial number of data points for evaluation of a small dataset in LOOCV [41, 42]. Therefore, LOOCV was used to compare the performance of our model on the ADNI and OASIS datasets with that of previous studies [43, 44, 45], and the results are reported here. Moreover, because the model’s performance can vary depending on random values, such as weight initialization, all experimental results were obtained with 20 different random seeds. Following the original paper [28], we set the teleport probability of APPNP, *α*, as 0.1. The MLP in Eq. 2 was structured as follows: size of hidden units = 64, embedding dimension *H* = 64, and the number of hidden layers *L* = 3 ^1^. Non-imaging demographic measures for both the ADNI and OASIS datasets included age, sex, and APOE, as illustrated in Table 1.

Four evaluation metrics used throughout this paper are defined as follows:

- 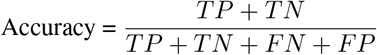
- 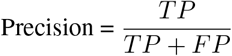
- 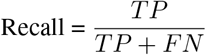
- 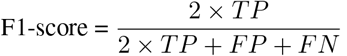

where TP, TN, FN, and FP denote true positive, true negative, false negative, and false positive, respectively.

We use accuracy as the main metric, as both datasets were well balanced.

**Table 1:** Non-imaging demographic measures for ADNI and OASIS datasets.

|  | ADNI | OASIS |
| --- | --- | --- |
| Age | 0-69, 70-79, 80+ | 0-59, 60-69, 70-79, 80+ |
| Sex | Male, Female | Male, Female |
| Gene | APOE4: 0, 1, 2 | APOE: 22, 23, 24, 33, 34, 44 |

### 3.2 Influence of hop sizes

As previously mentioned, over-smoothing has been regarded as the cause of performance deterioration in GCN. After multiple layers of convolution, the representations of distinctive nodes become indistinguishable. On the other hand, APPNP, which preserves the input information, is known to relieve this issue. Therefore, we trained and evaluated both the APPNP and GCN models by changing the hop size to report its influence on the overall performance, as a different number of propagation steps will result in distinct performance. The proposed GNN framework learns the neighboring nodes *K*-hops away from each node, and *K* ⊆{2, 4, 6, 8, 10}. The plots depicting the accuracy across all seeds and folds on each dataset are shown in Fig. 4.

**Figure 4:**
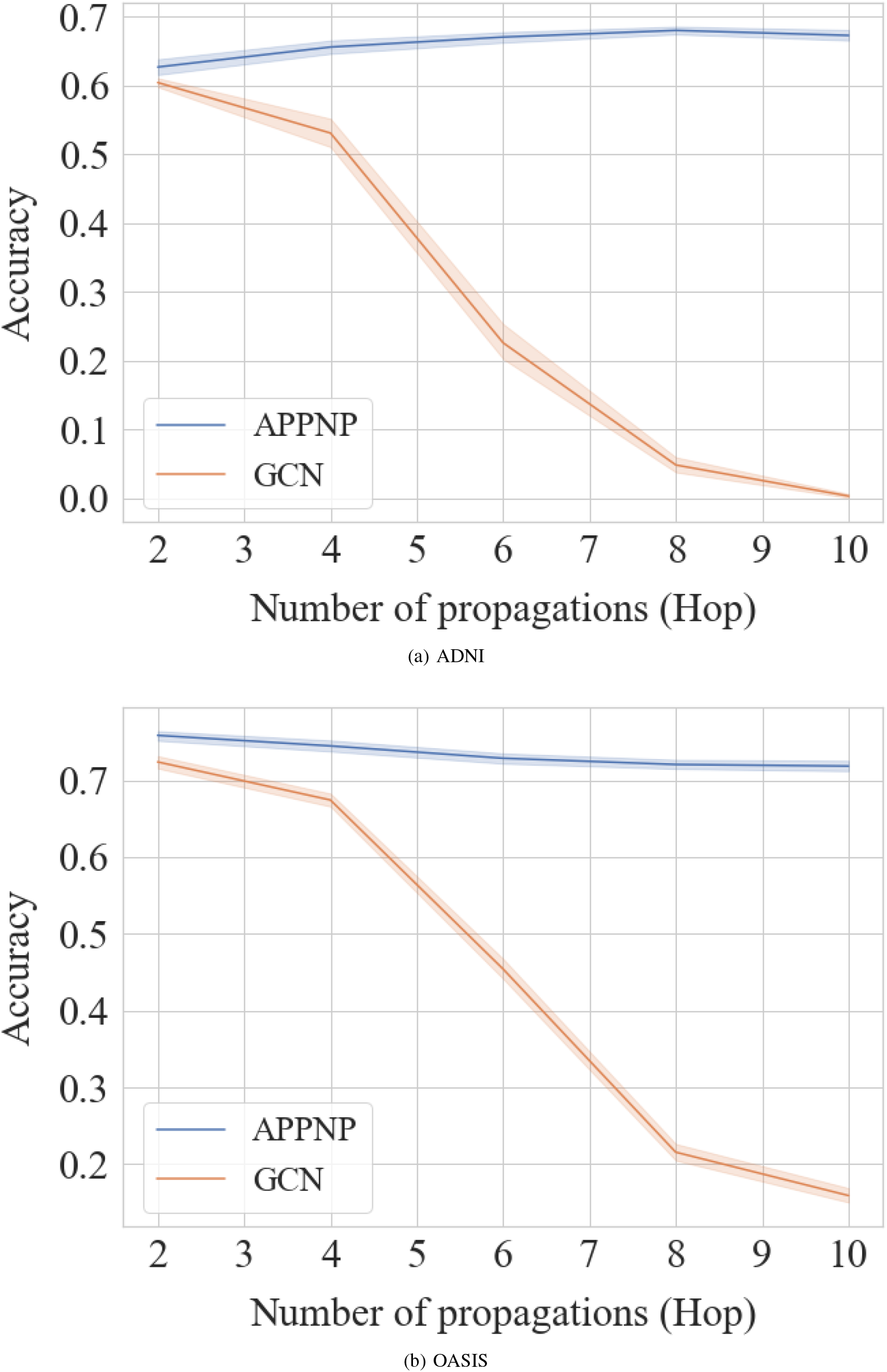
Classification results depending on different hop sizes. The shades represent the minimum and maximum values.

The best performance with an accuracy of 68% was obtained with the ADNI data when *K* = 8. A general tendency was observed, wherein the accuracy increased as *K* approached 8 and decreased thereafter. Moreover, the proposed APPNP model showed the best performance with the OASIS data when *K* = 2. Therefore, we used 8 and 2-hops, which is the optimal degree of K in the APPNP model, in the remainder of the experiments.

Interestingly, our model showed negligible variances depending on the hop size, whereas GCN showed an initial performance of 60%, which continuously decreased until it reached almost 0%. The accuracy drastically changed when the hop size ranged from 4 to 6 for ADNI and 4 to 8 for the OASIS data. This clearly demonstrates that the GCN model lacks the ability to utilize a higher number of propagation steps. For both datasets, the GCN model failed to learn effectively with hops over 4.

### 3.3 Influence of demographic measures

Because graph-based frameworks distinguish and learn different graph structures, incorporating an effective representation of such structures is necessary. Therefore, we further evaluated the performance of our proposed model by changing the combination of non-imaging demographic information such as age, sex, and APOE genotype. The results are shown in Fig.5 with multiple initialization seeds for convergence. The first observation showed approximately 10% difference between the worst and the best-performing demographic measures for the ADNI and OASIS data. In other words, incorporating a poor combination of demographic measures can significantly decrease the performance of the model.

With the ADNI data, when considering only one type of non-imaging demographic information, we observed that using the data on sex yielded the highest accuracy, whereas age showed the poorest accuracy. We also found a general tendency of increasing performance when the APOE genotype was combined with other demographic measures. As shown in Fig. 5, “APOE genotype, Sex” and “APOE genotype, Age” yielded better performances than “Age, Sex.” The best performance was obtained with all three demographic measures.

**Figure 5:**
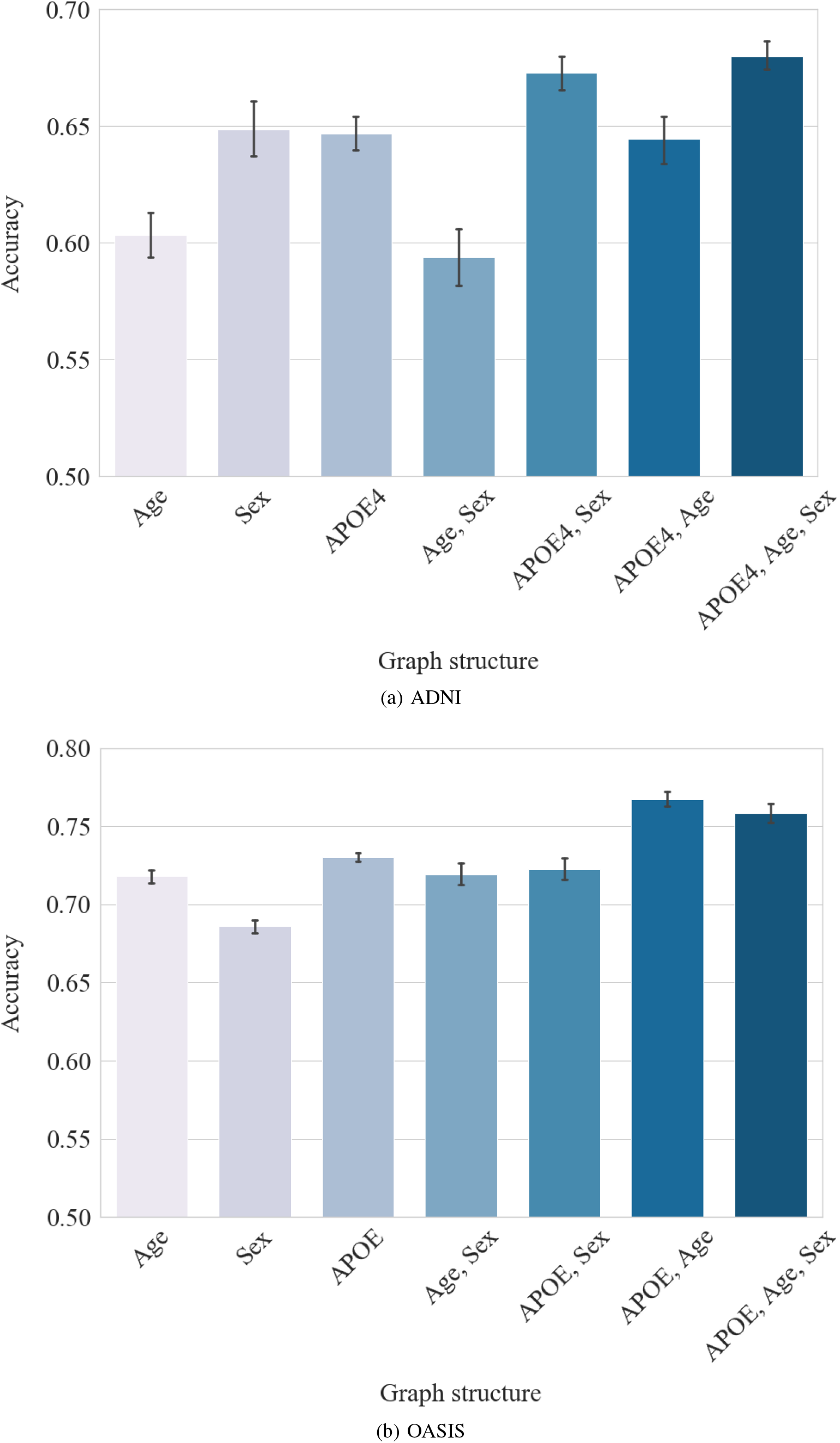
Influence of the demographic measures on the classification performance. The barplot illustrate the average accuracy across all seeds.

With the OASIS data, using the APOE genotype information showed the best performance among “Age,” “Sex,” and “APOE genotype,” as shown in Fig. 5, which was similar to the results of ADNI. However, because “Age” showed better performance than “Sex,” “Age, Sex” showed comparable performance to “APOE genotype, Sex.” Furthermore, the “APOE genotype, Age” combination was better than that of “APOE genotype, Sex.” Generally, our model showed increased performance when incorporating the APOE genotype information in the graph. Similarly, as in the ADNI data, “APOE genotype, Age, Sex” showed the second-best performance.

For uniformity, we used the combination of “APOE genotype, Age, Sex” for both ADNI and OASIS as the default graph structure in this study.

### 3.4 Comparison with other methods

We further compared our classification results with several other well-defined classifiers: K-nearest neighbor (KNN), MLP, linear regression, convolutional neural networks (CNN), and GCN.

With the ADNI dataset, KNN demonstrated the worst classification performance from 35% to 55% in terms of all four measures, and CNN slightly outperformed KNN. The superior performance of our proposed APPNP framework when compared with MLP indicates that incorporating non-imaging information in a graph structure improves the performance. Furthermore, the best performance is repeatedly observed with our APPNP model, which is approximately 7% higher than that of the conventional GCN (Table 2).

**Table 2:** Summarized comparison results with ADNI in terms of four evaluation metrics.

|  | Accuracy | Precision | Recall | F1-score |
| --- | --- | --- | --- | --- |
| KNN | 53.22 | 55.0 | 35.48 | 43.13 |
| Linear | $62.17 \pm 1.84$ | $62.29 \pm 2.16$ | $61.93 \pm 2.24$ | $62.08 \pm 1.66$ |
| CNN | $59.27 \pm 4.21$ | $59.16 \pm 4.41$ | $60.8 \pm 5.36$ | $59.85 \pm 4.08$ |
| MLP | $62.82 \pm 1.43$ | $63.05 \pm 1.41$ | $61.93 \pm 2.68$ | $62.46 \pm 1.79$ |
| GCN | $60.4 \pm 1.61$ | $61.09 \pm 1.66$ | $57.25 \pm 2.31$ | $59.1 \pm 1.86$ |
| Homogeneous APPNP | $66.04 \pm 1.1$ | $67.77 \pm 1.62$ | $61.29 \pm 0.0$ | $64.36 \pm 0.74$ |
| **APPNP** | **$67.98 \pm 1.41$** | **$69.34 \pm 1.66$** | **$64.51 \pm 2.09$** | **$66.82 \pm 1.54$** |

KNN showed performance improvement of approximately 60% with the OASIS dataset when compared with the ADNI dataset; however, this value was again the lowest. Similar to the results for ADNI, a general tendency was observed where the linear regression model performed better than CNN. CNN outperformed linear regression in only one of the 4 measures, i.e., recall, which was negligible because the standard deviation was different. Importantly, our model showed the best performance of approximately 75% for all four evaluation measures (Table 3).

**Table 3:** Summarized comparison results with OASIS in terms of four evaluation metrics.

|  | Accuracy | Precision | Recall | F1-score |
| --- | --- | --- | --- | --- |
| KNN | 60.0 | 60.16 | 59.16 | 59.66 |
| Linear | $69.39 \pm 0.65$ | $72.23 \pm 0.65$ | $63.0 \pm 1.09$ | $67.3 \pm 0.82$ |
| CNN | $64.58 \pm 2.52$ | $64.85 \pm 2.61$ | $63.75 \pm 3.1$ | $64.27 \pm 2.57$ |
| MLP | $70.66 \pm 0.51$ | $73.53 \pm 0.64$ | $64.58 \pm 0.68$ | $68.76 \pm 0.56$ |
| GCN | $72.37 \pm 1.91$ | $72.0 \pm 1.92$ | $73.24 \pm 2.49$ | $72.6 \pm 1.98$ |
| Homogeneous APPNP | $68.02 \pm 2.84$ | $67.87 \pm 2.93$ | $68.5 \pm 3.05$ | $68.17 \pm 2.81$ |
| **APPNP** | **$75.83 \pm 1.41$** | **$75.77 \pm 1.28$** | **$75.95 \pm 2.52$** | **$75.84 \pm 1.61$** |

### 3.5 Effectiveness of heterogeneous APPNP

In addition to comparison with baseline classification methods, we tested the effectiveness of the proposed APPNP model. While our model can facilitate the utilization of distinct *heterogeneous* nodes (i.e., different types of nodes) through the *subject–demographics graph*, *homogeneous* APPNP uses a *subject–subject graph* that connects the subjects’ nodes if they share commonalities among the demographic measures. Demographic nodes are absent from the graph in this model. We compared the performance of both models, and the results are illustrated in Tables 2 and 3.

The positive effect of incorporating the demographic measures in a heterogeneous manner with the *subject– demographics graph* was observed in this comparison study. Our heterogeneous APPNP model continuously showed superior performance when compared with the homogeneous model for both the ADNI and OASIS datasets. In addition, the homogeneous model was inferior to the MLP model for the OASIS dataset. The overall differences in performance between these two methods are approximately 2% and 7% for the respective data. We reason that the homogeneous model oversimplified the graph structure and failed to account for the complex relations among the subjects, especially for the larger graph.

## 4 Conclusion

In this paper, we proposed a novel framework for classifying AD using a graph neural network by incorporating heterogeneous data and demonstrated that it outperforms the conventional deep learning approaches. Our proposed method utilizes the *subject–demographics graph* to integrate the resting-state functional connectivity and demographics, whereas previous studies manually defined the similarities among the nodes to connect them through the edges. We expect that the combination of imaging and non-imaging features by constructing a bipartite graph would improve the generalizability. Another advantage of this deep learning framework is that it automatically learns the essential features for classification in an end-to-end manner. In other words, it does not rely on handcrafted features, which were common in previous studies.

APPNP showed superior and robust performance than the conventional GCN, even when the number of propagations increased. Although there is limited research on explaining why the majority of GCN models are shallow, several studies have empirically shown that these architectures are generally applicable for 2 to 3 layers [46, 25, 47]. As shown in Fig. 4, GCN shows a drastic performance drop after 4 hops, both with the ADNI and OASIS datasets. On the other hand, APPNP rigorously demonstrated better performance than GCN as the size of hops increased, because it applied personalized PageRank to tackle the over-smoothing issue. The ability to stack multiple propagation steps resulted in larger receptive fields to gather more information for learning. Consequently, the classification accuracy of APPNP was always higher than that of GCN.

Translating heterogeneous data into a graph structure plays an essential role in the classification performance. For neuroimaging data, we used static RSFC as one of the features in the graph. Utilizing dynamic functional connectivity within the graph could be beneficial because it reflects time-varying dynamic behavior [48, 49]. However, because there is no current consensus on the methods to calculate the dynamic functional connectivity, it needs to be carefully examined. In terms of non-imaging information, integrating different types of demographic measures in a graph significantly affects the classification accuracy, as shown in Fig. 5. For instance, solely using the APOE genotype information resulted in better performance than the combination of “Age, Sex” for both the ADNI and OASIS data. This can be attributed to the identification of a highly correlated relationship between the APOE genotype and AD onset [50, 51]. Within the ADNI dataset, “APOE genotype, Sex” showed better performance than “APOE genotype,” which may reflect a difference in vulnerability to the disease depending on sex [52, 53]. On the other hand, decreased accuracy value of the “APOE genotype, Age” compared with “APOE genotype” might be caused by limited data representation.

Our experiments also demonstrated that utilizing heterogeneous data through a *subject–demographics graph* in APPNP was critical for the classification accuracy. As depicted in Tables 2 and 3, the results showed higher accuracy with the ADNI dataset than the homogenous APPNP. A similar tendency was observed with the OASIS data. Integrating heterogeneous data in a *subject–demographics graph* is not only convenient for adding features without considering the importance weights, but also increases the representation power. Moreover, it is able to handle missing features that are common in the dataset. Other graph construction strategies can be applied to combine neuroimaging and diverse non-imaging demographic data, such as hippocampal volume, cortical thickness, and biochemical examination results. One could consider multiple graph structures, such as building an adjacency matrix per demographic measure. Furthermore, important non-imaging features can be filtered from all information through self-attention weight learning. These sophisticated approaches can help identify the important features of dissipated information and improve the classification results.

Due to its convenience, a binary classification of CU and AD patients was performed in this study. However, because AD lies in the spectrum, future studies are necessary to expand our proposed method by performing multi-class classification as well as predicting the prognosis. This can be done by optimizing parameters such as the number of hidden layers, feature vector size, and loss function used in this study. In addition, we utilized two balanced datasets to test the applicability of the proposed model. However, substantial imbalance among classes can be found in certain types of data. Therefore, in future studies, we would like to explore how graph neural networks can be used for large-scale imbalanced datasets.

## Footnotes

1 Without loss of generality, the size of the hidden units, embedding dimension, and number of hidden layers can be freely determined with different values. Empirically, it was found that using shallow and thin networks is suitable for a small dataset, as in this study, by preventing overfitting.

